# H5N5 virus transmission is driven by repeated cross-Atlantic introductions and transmission among high latitude seabirds

**DOI:** 10.64898/2026.10.29.762228

**Authors:** Clara R. Malekshahi, Nicole St. Clair Knobloch, Alvin Crespo-Bellido, Alexa Simulynas, Stephanie Goya, Leigh Papalambros, William J. Fitzsimmons, Alexander L. Greninger, Adam S. Lauring, Jonathan Runstadler, Martha I. Nelson, Wendy Puryear, Louise H. Moncla

## Abstract

In November 2025, the first global H5N5 infection was reported in a person in Washington State. Previously, these viruses had only been detected in seabirds and shorebirds near the Atlantic Ocean, suggesting potentially distinct ecology from dominant H5N1 strains. Using 354 global H5N5 genomes, we show that H5N5 viruses have been carried by high latitude seabirds and shorebirds across the Atlantic at least 9 times, with introductions more frequent in the fall and spring. We reconstruct a plausible path of cross-continental movement via Arctic or sub-Arctic regions, followed by a year of cryptic circulation prior to human spillover. These data show that H5N5 viruses have undergone far more intercontinental movements than previously recognized, and exhibit host and geographic patterns that are fundamentally distinct from dominant H5N1 lineages. The distinct host ecology, geography, and movement, paired with the capacity to infect a person, suggest that enhanced surveillance may be warranted.

## Introduction

In 2021, highly pathogenic avian influenza H5N1 viruses of the 2.3.4.4b clade were introduced into North America, sparking a panzootic that has resulted in mass mortality in wild birds, the culling of hundreds of millions of domestic poultry, and frequent spillovers into wild and domestic mammals (1–3). These viruses continue to circulate among wild birds in North America, driven by transmission in wild Anseriformes (ducks, geese, and swans) (4), with periodic introductions of new H5N1 lineages from Europe and Asia (4,5). While H5N1 viruses have been the predominant subtype in the Americas, additional introductions of reassortant, clade 2.3.4.4b H5N5 viruses have also occurred (5,6). This subset of H5N5 viruses emerged in eastern Europe in 2020/2021 following reassortment with local low pathogenicity avian influenza viruses (7). There are 3 documented introductions of these viruses from Europe to North America, which are collectively classified as “A6” genotype viruses (5,6,8). A6 H5N5 viruses have been detected far less frequently than H5N1 viruses, with H5N5 cases localized to the Northwest Atlantic coast in gulls and related seabirds, particularly Great black-backed gulls (*Larus marinus*) and Northern fulmars (*Fulmarus glacialis*) (5). Like H5N1, however, H5N5 viruses have been detected in scavenger birds, foxes, seals, and raccoons, suggesting the potential risk for mammalian infection and adaptation (5,6).

In November 2025, a person in Washington state, U.S.A., was hospitalized with an HPAI H5N5 virus, marking the first human infection with an H5N5 virus globally (9). Genomic characterization classified this virus into the A6 genotype (9), suggesting that it was most closely related to viruses detected among seabirds on the Atlantic Coast, over 5,000 kilometers away. Until then, HPAI H5N5 viruses had not been detected in the Pacific flyway, suggesting that A6 virus ecology may be distinct from dominant H5N1 strains.

To reconstruct patterns of viral introduction and spread within North America, we curated a global dataset of 354 HPAI H5N5 viral sequences and used Bayesian phylodynamic approaches to model transmission across hosts and geographic space. We identify repeated transmission events between Europe and North America, likely facilitated by seabird transmission at high latitudes. We trace a plausible path of transmission from the Atlantic to Pacific via the Arctic circle, followed by cryptic transmission of at least one year prior to the human infection. In contrast with the extensive spread of HPAI H5N1 viruses by Anseriformes (ducks and geese) across North America, our data suggest that H5N5 viral ecology and movement is uniquely facilitated by high-latitude seabirds. These data collectively suggest that surveillance strategies may need to be tailored to account for the distinct ecologies supported by co-circulating HPAI lineages. These data, combined with sporadic detections in mammals in the Pacific, suggest that H5N5 viruses are capable of spillover, but likely heavily under-surveilled.

## Results

### Surveillance and detection of H5N5 viruses among Northeastern seabirds

Since the first detection of H5N5 in Massachusetts in Great black-backed gulls (*Larus marinus*) in January 2023, ongoing surveillance of wild birds in the northeastern United States has continuously detected low prevalence HPAI subtype H5N5 A6, primarily in seabirds, with additional repeated detections in birds of prey and scavengers. These H5N5 A6 viruses were consistently detected even as H5N1 genotypes emerged and disappeared within the same region, suggesting that these H5N5 lineages persisted and co-circulated alongside predominant H5N1 strains. The majority of H5N5 detections occurred in Great black-backed gulls (n = 17), with additional positives in Herring gulls (n = 2), seaducks (1 each of a Common Eider, Red-breasted Merganser, and Bufflehead), a shorebird (1 Sanderling), and from birds of prey and scavenger species: Red-tailed hawks (n = 2), Common ravens (n = 2), a Bald eagle (n = 1), Black vulture (n = 1), and Turkey vulture (n = 1). In total, genetic sequencing confirmed the presence of 31 HPAI H5N5 A6 wild bird cases in Maine and Massachusetts over the study period, with 26 producing sequences suitable for subsequent use. Among birds testing positive, over 80% were found deceased. Of those birds that were admitted to a rehabilitation facility, all exhibited neurological clinical signs including lethargy, astasia, and torticollis and subsequently died or were humanely euthanized.

We combined the 26 sequences from our study site with all globally available H5N5 A6 full genome virus sequences (see Methods for details) (10,11), resulting in a final dataset of 354 viral genomes. These data represent all available sequences from infections sampled from November 2021 to May 2026 across 15 countries and 52 host species (Fig. 1, Supplementary Figure 1). Most sequences represent samples collected from Charadriiformes, the avian order that includes gulls and terns, with the largest representation from Great black-backed gulls. Large numbers of sequences also came from Northern fulmars (order: Procellariiformes), corvids (order: Passeriformes), and eagles (order: Acciptriformes) (Fig. 1B). Global genomes therefore align with the distribution of host species samples from our study site.

**Figure 1.**
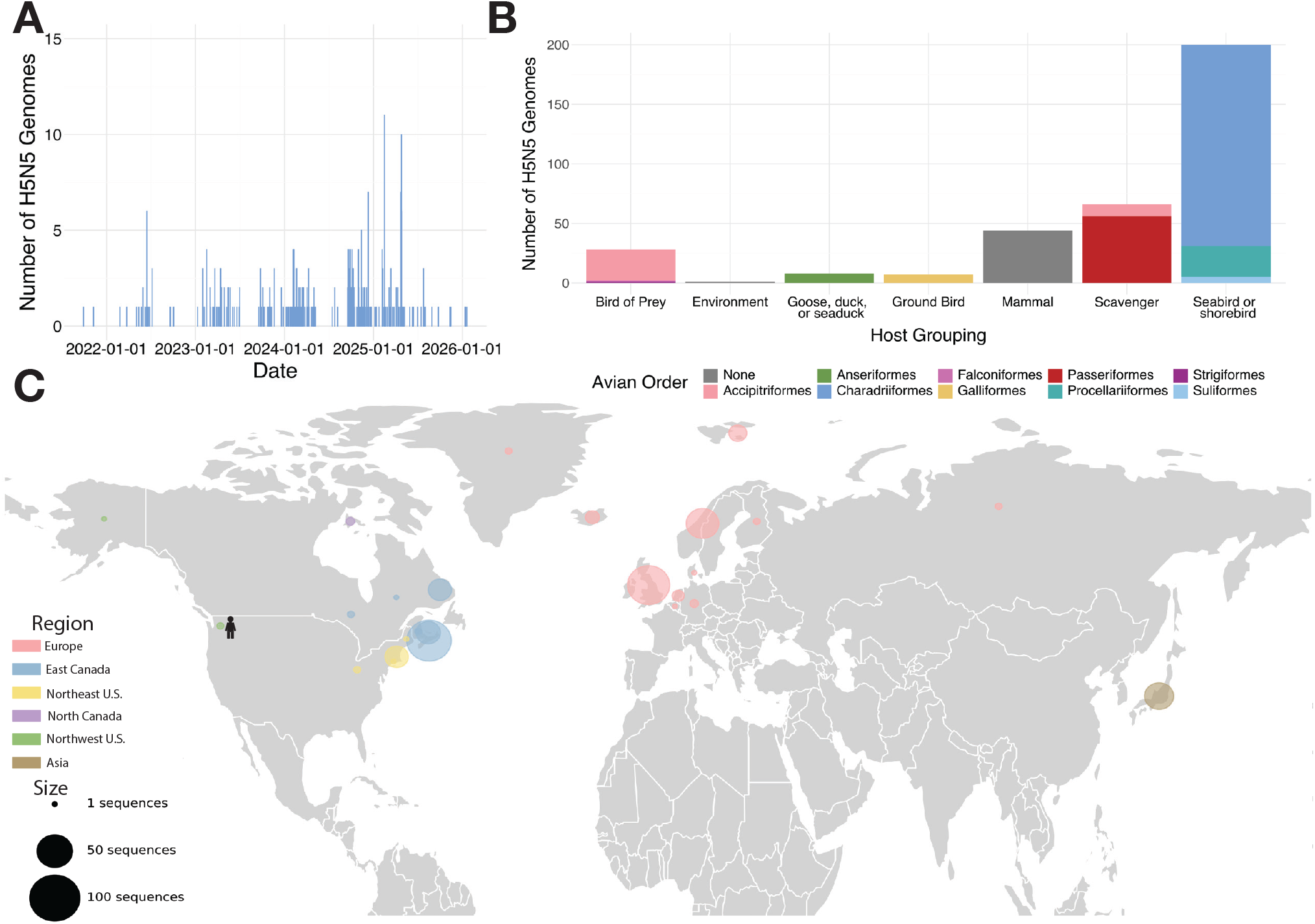
Sequences are distributed across time, location, and host groups. We collated 354 concatenated H5N5 A6 genomes, sequenced from 2021 to 2026 **(A).** Concatenated genomes are distributed across ecologically-defined host group (birds of prey; geese, ducks, or seaducks; ground birds; mammals; scavengers; seabirds and shorebirds), colored by the avian taxonomic order **(B)**. Concatenated genomes are distributed globally, where size indicates the number of sequences from a country (Europe, Asia), state (United States), or province (Canada), and color indicates the region assigned for future analysis **(C)**.

### Frequent cross-continental movement of H5N5 viruses is concentrated in spring and fall

Early cross-continental introductions of clade 2.3.4.4b viruses have been hypothesized to originate from migratory seabirds (1,5). To determine the frequency of viral introductions between continents, we grouped sequences by the region of sampling, generated concatenated full genome alignments, and inferred Bayesian phylogenetic trees with geographic diffusion modeled as a discrete trait (see Methods for details). We inferred tree topologies using three potential demographic models and evaluated model fit by estimating the marginal likelihood of each model using Stepping Stone sampling (12). An exponential growth model was selected as the best fit and used for all subsequent reconstructions (Supplementary Table 1, see Methods for details).

All sampled North American H5N5 A6 viruses descend from viral diversity circulating in Europe, consistent with prior studies (5,6) (Fig. 2A). Though currently classified as a single genotype, we infer that H5N5 A6 viruses have been introduced to North America 9 to 10 (mean

**Figure 2.**
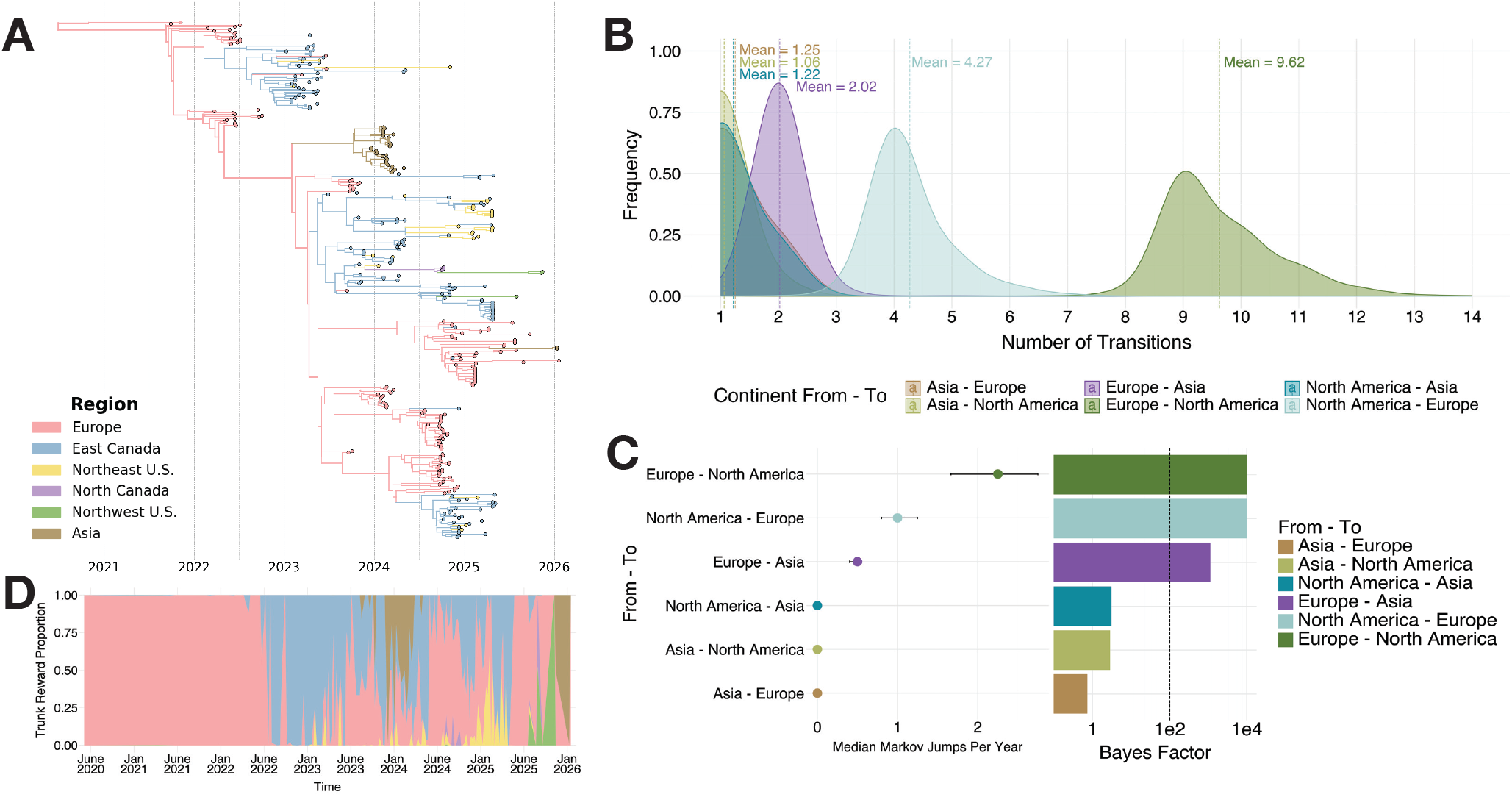
The H5N5 A6 genotype is comprised of multiple introductions from Europe to North America. We visualize the Maximum Clade Credibility tree of 354 concatenated A6 H5N5 viral genomes, with region inferred across the phylogeny as a discrete trait and represented through color **(A).** For each pairwise transition between continents, we infer the history of Markov Jumps and plot the frequency of inferred jumps across the posterior set of phylodynamic trees (mean shown as a dashed line, Bayes factors shown in Panel C) **(B)**. For each pairwise transition between continents, we calculate the Bayes Factor support (logged) and plot the median number of Markov Jumps (transitions) per year (95% HPD of Markov Jumps across the posterior set of trees shown as error bars) **(C)**. We infer the trunk reward proportion of the Maximum Clade Credibility tree in each region using Markov Rewards, averaged per week, where color represents the region **(D)**.

= 9.62, 95% HPD = 8-11) independent times (Fig. 2B, Supplementary Figure 2), including three introductions previously described in Rahman et al. (2024) (5). All North American introductions occurred from Europe into eastern Canada, with three introductions leading to subsequent flow southward to the northeastern United States (Fig. 2A). We find strong support (Bayes Factor >100) for bi-directional transmission between Europe and North America, with small but well-supported transmission events inferred from North America to Greenland and Scandinavia (Fig. 2C). We also reconstruct two introductions of an H5N5 A6 virus from Europe into Asia, one of which resulted in a documented outbreak in crows in Japan in 2023-24 (5,13). Roughly 50% of inferred Markov rewards (a metric for transmission) was maintained in both Europe and eastern Canada from 2020 through 2023, supporting these regions as key sources of viral persistence (Fig. 2D, Supplementary Figure 3).

Global spread of clade 2.3.4.4b viruses has been driven by wild, migratory birds, which exhibit seasonal variation in movement (2). To determine whether H5N5 A6 virus introduction patterns varied across seasons, we calculated the cumulative number of transitions between North America and Europe over time (Fig. 3A). Transitions from Europe to North America have occurred repeatedly since 2021, with the highest estimated transitions occurring in March, April, May, June, October, and November (Fig. 3B). While the inferred number of cross-continental transitions is consistent across the posterior, mapping to specific months was variable, precluding robust statistical analysis of month to month transition rates (Supplementary Figure 4, 5). Although the timing of cross-continental introductions between Europe and North America is difficult to accurately estimate, we infer that H5N5 A6 viruses represent at least 9 to 10 introductions into North America, and that cross-continental movement may be most common during the spring and fall. These data suggest that H5N5 viruses experience cross-continental movements at higher frequencies than previously recognized.

**Figure 3.**
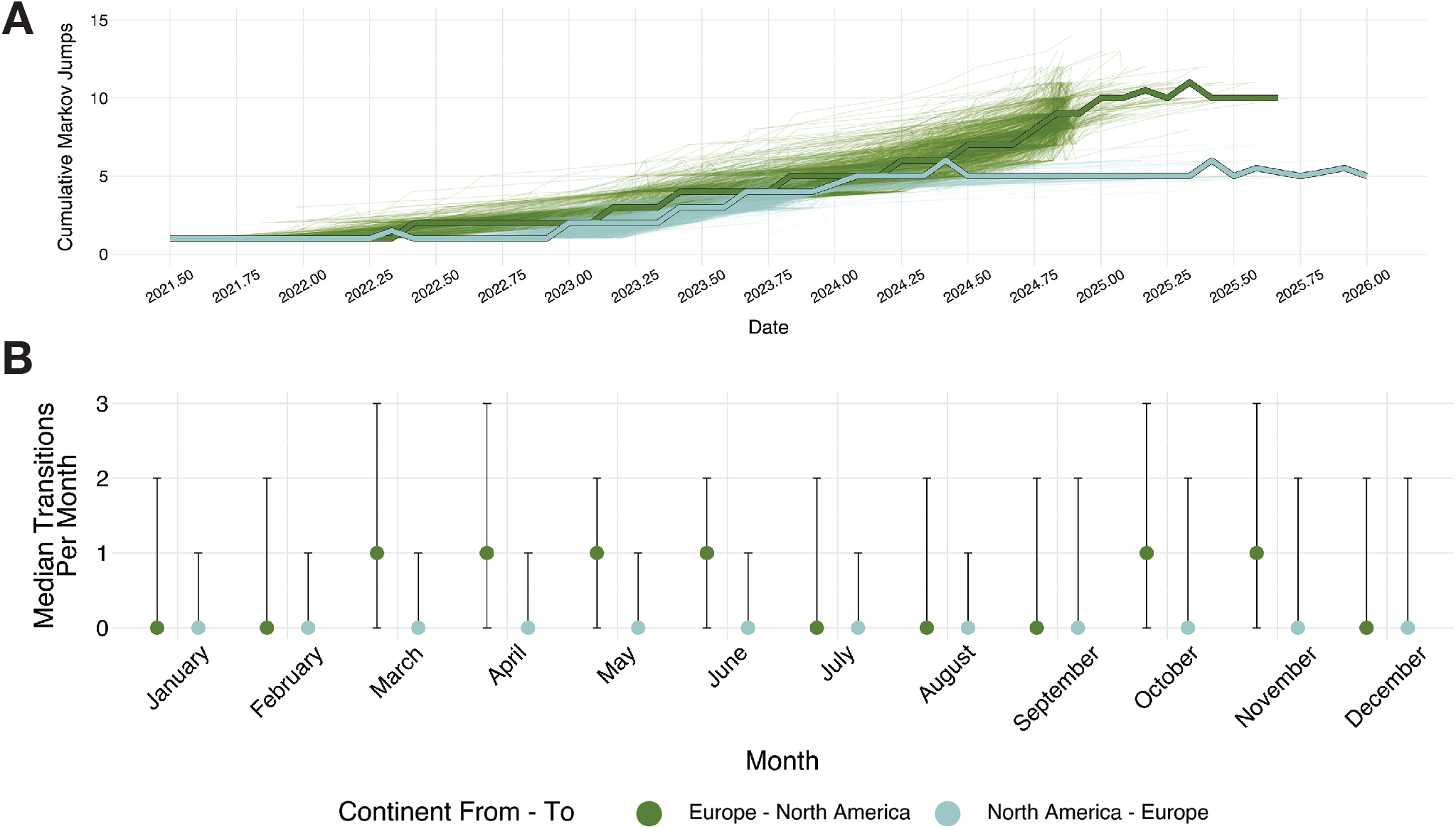
Cross-continental movement between Europe and North America exhibits seasonality. We inferred transitions between Europe and North America as Markov Jumps and calculated the cumulative number of transitions over time across the posterior set of trees (median in bold) **(A).** We also calculated the median number of transitions occurring in each month between Europe and North America across the posterior distribution of trees (error bar = 95% HPD) **(B)**.

### H5N5 virus transmission is driven by seabirds across northern latitudes

While clade 2.3.4.4b H5N1 virus transmission is mostly driven by ducks and geese (order: Anseriformes) (4), H5N5 viruses have been primarily detected in gulls and other seabirds (5) (Fig. 1B). We next modeled host groups as a discrete trait (see Methods for details), and inferred viral dispersal using a Bayesian continuous phylogeographic model. While H5N1 viruses cause recurrent, large outbreaks in domestic poultry, A6 viruses have only been detected in 7 out of 354 total Galliformes (turkeys, pheasants, and chickens) in our dataset, suggesting distinct patterns of spread. With the exception of a pair of pheasant sequences isolated from England, these Galliformes sequences are phylogenetically unrelated and represented isolated events, implying that these viruses rarely cause detectable spillovers in Europe and North America (Supplementary Figure 6A). Within the H5N5 phylogeny, all introductions into North America were inferred to have a seabird host origin, with sporadic transmission into scavengers (crows and vultures), birds of prey, and nonhuman mammals (Fig. 4A, Supplementary Figure 6A). Transmission within Europe and North America is also associated with seabirds, which were inferred as the predominant source host across all continents (Supplementary Figure 6). Viral transmission was restricted to high latitudes, with most detections clustered to Northern Atlantic coasts north of 35.9°N (Fig. 4B, Supplementary Figure 7). Transmission within eastern and northern Europe occurs primarily in high-latitude, coastal countries, with frequent cross-Atlantic movements into eastern Canada that then spread along the coast to the northeastern United States (Fig. 4B, Supplementary Figure 7). Diffusion rates (a measure of dissemination speed) (14) varied little across North America (Fig. 4C), and were only slightly more variable within Europe (Fig. 4C), suggesting that these viruses spread similarly across locations. The majority of viral lineages introduced into North America only spread a limited distance (<1,000 km) from the point of introduction (Fig. 4D), supporting localization to coastal states. Overall, these data suggest that H5N5 virus transmission is primarily mediated by seabirds, restricted to high latitudes, and very rarely causes detectable spillovers to agriculture, marking patterns that are entirely distinct from dominant H5N1 strains.

**Figure 4.**
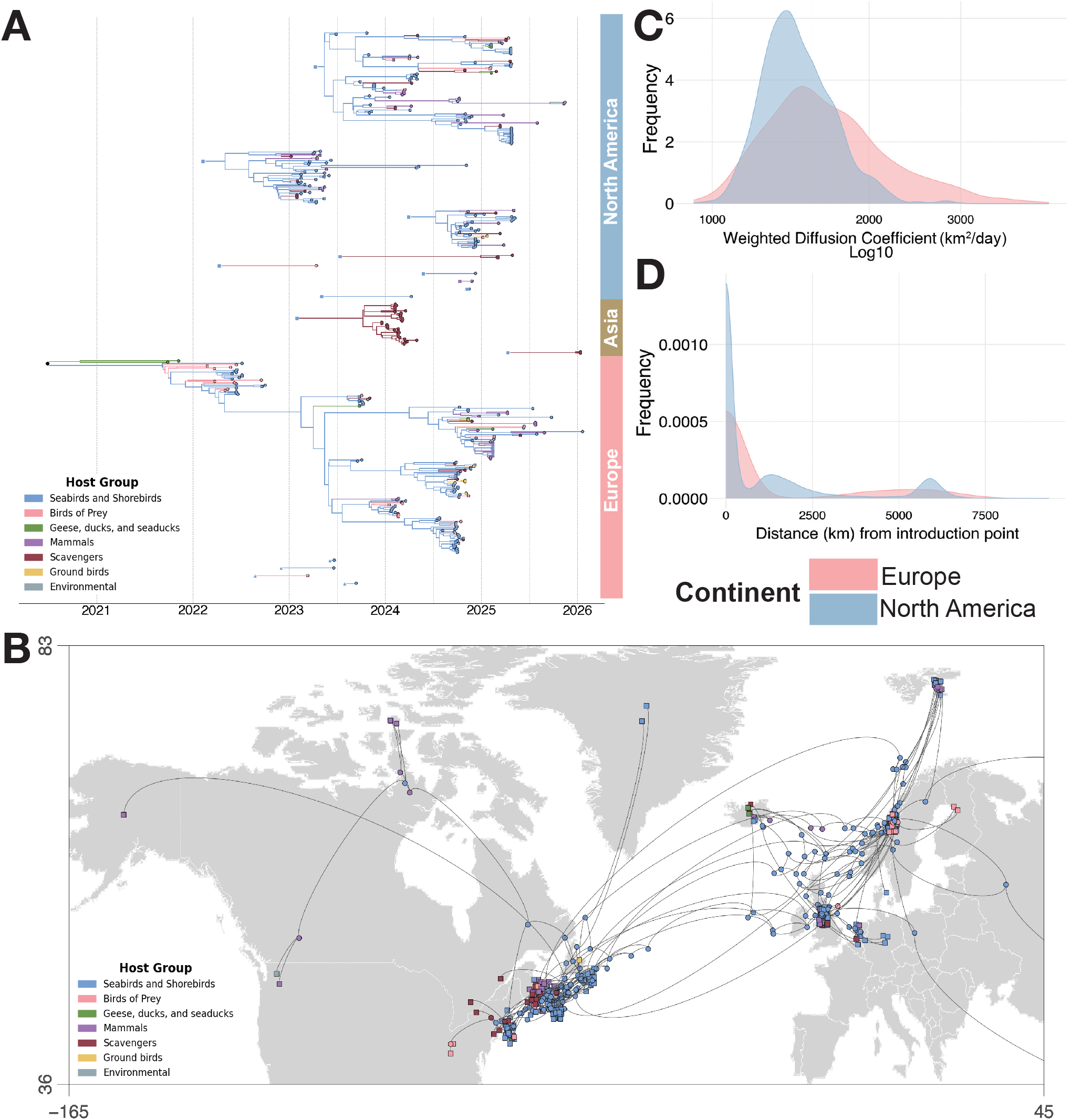
Global H5N5 transmission is driven by seabirds across high latitudes. We visualize the Maximum Clade Credibility tree of 354 concatenated genomes, with host group inferred across the phylogeny and represented as color, exploded by cross-continental introductions, with origin continent indicated by shape (square = Europe, triangle = North America), and destination continent indicated by colored bars at right **(A).** We visualize the Maximum Clade Credibility tree on a geographic map of Europe and North America, showing the movement of inferred (circles) and sequenced (squares) HPAI H5N5 A6 genomes, colored by host group (not shown: sequences from Russia and Japan) **(B)**. We calculated the weighted diffusion coefficient of viral lineages starting and ending in Europe or North America across the posterior set of trees (not pictured: Asia) **(C)**. We calculated the maximum distance traveled (kilometers) from the first node in each introduction to the continent across the posterior set of trees (not pictured: Asia) **(D)**.

### Cryptic transmission and possible cross-Arctic spread occurred for a year prior to human spillover

In November 2025, a person was infected with an H5N5 virus in Washington State, USA, nearly 5,000 kilometers from the Atlantic Coast (9) where most detections have been localized (5,6) . Case investigation found that this person was likely exposed via interaction with backyard ducks, from which HPAI was identified in samples from an apparently healthy duck and backyard sediment (9). Although these viruses had not previously been detected in any birds within the Pacific Flyway, case investigation concluded that the most likely route of exposure was from introduction via wild bird into the backyard environment (9), raising questions about how these viruses traversed the continent prior to spillover. To determine a plausible path by which these viruses spread to the Pacific, we included the human sequence in our phylogenetic reconstruction, along with a sequence later sampled from a polar bear near Alaska. The viral sequence isolated from the human case is most closely related to an environmental sample from Washington collected around the same time and location, which was presumably sampled from the backyard sediment (November 14, 2025). Both sequences then nest most closely with sequences collected from ringed seals and a Northern fulmar in Nunavut, Canada in October 2024. These Nunavut detections descend from an introduction into eastern Canada in April 2023 (95% HPD = February 2023 - May 2023) that we infer circulated on the East Coast and within northern Canada for roughly 1.5 years before the infection in Washington state (Fig. 4B, Fig. 2D). These data suggest possible dissemination from the east to west coast via movement across the Arctic circle, a breeding ground for Northern fulmars, kittiwakes, and a variety of gulls, and a route of westward migration for jaegers, Arctic terns, and eiders (15,16). The branches connecting the human-derived sequence to its closest ancestors (the ringed seals in northern Canada) span roughly 1.2 years, indicating substantial unsampled diversity separating these cases (Table 1). The only other sample from the northwestern United States was isolated from a polar bear sampled in Alaska in August 2025, and exhibits a similar branch length of just under one year (Table 1). Given a clock rate that is consistent with continuous transmission, these data are compatible with cryptic transmission in birds for at least one year prior to the human infection (Supplementary Figure 5). These long branches represent outliers, with most introductions of H5N5 viruses remaining within short distances of their introduction point in both North America and Europe (Fig. 4D). Surveillance within high latitudes in northern Canada are sparse, and viruses circulating here may be undersampled, providing a plausible explanation for this long period between detections. Together, these data suggest that these viruses may have been circulating cryptically in wild birds for at least a year prior to spillover, and highlight the capacity for these viruses to be spread coast to coast via Arctic latitudes.

**Table 1.** Long-term cryptic circulation of branches crossing North America. We inferred dates and branch lengths (mean, 95% HPD) of the three genomes from the northwestern United States, as well as the inferred ancestor of the human infection and the related environmental sequence, across the posterior set of trees.

| Object | Strain Name | Date | Mean branch length (yrs) |
|---|---|---|---|
| Internal Node | None | 2025.713 | 0.990 [0.662 - 1.247] |
| Sequence | A/Washington/2148/2025 | 2025.858 | 0.196 [0.023 - 0.481] |
| Sequence | A/environment/Washington/04434-001-v/2025 | 2025.868 | 0.207 [0.034 - 0.492] |
| Sequence | A/Polar Bear/AK/26G03248/2025 | 2025.581 | 0.986 [0.851 - 1.168] |

## Discussion

This work highlights the complexity of H5 virus ecology within the wild bird reservoir. We find that HPAI H5N5 A6 viruses exhibit a unique ecology dominated by transmission among seabirds at high latitudes, and are frequently moved between Europe and North America.

Though most introductions remained localized to the Atlantic coast, one lineage travelled from the Atlantic to the Pacific coasts via the Arctic circle, circulating for at least one year prior to a human spillover. Our data show that ecology and transmission of H5N5 viruses is wholly distinct from that of dominant H5N1 lineages, which are maintained by Anseriformes and circulate throughout the continent. While current surveillance systems are designed to capture H5N1 viruses, our data suggest that gaps exist in our understanding of H5N5 reintroductions, continued circulation, and viral spread that could be better targeted by surveillance.

We find a high degree of circulation within the northern Atlantic region and frequent cross-continental spread, with some evidence for enhanced cross-continental movements in the fall and spring. While limited sequence numbers precluded robust estimation of monthly introduction rates, these observations align with known avian migratory movement patterns that could facilitate spread. Fall and spring introductions could be facilitated by short- and medium-distance migrations of gulls and other seabirds to and from breeding colonies in the United Kingdom, Scandinavia, Iceland, Greenland, and eastern Canada (1,5). In particular, Iceland has been previously identified as a “stepping stone” for HPAI H5N1 migration from Europe to North America in this manner (17). Great black-backed gulls (*L. marinus*), in which detections are most common in both Europe and North America, are found in resident and medium-distance migrant populations in Greenland, Canada, and the United States, from New England down to the mid-Atlantic coast (18). Migration patterns across the northern Atlantic Ocean ultimately align with our finding that transmission of these viruses is primarily associated with seabirds of the Charadriiformes and Procellariiformes orders, and we speculate that migratory movements represent plausible explanations for the frequent cross-continental transmission we infer. Future, expanded studies focusing on surveillance within these species may enable more robust estimates of specific months when introduction risk is highest.

One surprising feature of A6 H5N5 viruses was the appearance of a human infection on the Pacific coast, without any prior detections in that region. While most inferred viral introductions remained close to the Atlantic coast, two spread over 5,000 kilometers, one of which ultimately led to the human infection in Washington state. We infer a plausible path in which these viruses arced up from the northeastern United States into Nunavut, Canada, and then back down to the Pacific Northwest. Although our phylogenetic reconstructions are limited by small sample sizes, data on the distribution of wild birds in these regions lend credibility to these inferred paths. Great black-backed gulls have been abundant on the Massachusetts coast since 1931, nesting on islands, in marshes, and on barrier dunes along the coast (19). These birds span northern latitudes in North America and Europe, and occasionally migrate along the Atlantic coast (18). Since HPAI H5N5 A6 viruses were first detected in New England in 2023, Northern fulmars have also been spotted in areas where Great black-backed gulls reside, including off the coast of Provincetown, Massachusetts, at Stellwagen Bank, a National Marine Life Sanctuary 25 miles east in the Northwest Atlantic Ocean, and on ship-based bird surveys in the Atlantic Ocean 100 miles east of Cape Cod (20–22). Northern fulmars (*F. glacialis*) are high-latitude seabirds that spend their lives on the open ocean in Arctic regions such as Alaska, Canada, and Greenland and winter as far south as the New England coastline (23,24). Within the Arctic, other gull species (e.g., Herring gulls) breed and migrate, but also maintain ranges along the Pacific coast (25). While current data is insufficient to map the exact route of spread, we speculate that H5N5 A6 viral movement could include cross-continental and trans-Arctic spread linked to transmission in Northern fulmars and gull species (e.g., Herring gulls). Unique behavioral traits may also contribute to the H5N5 virus transmission in these species. For example, Northern fulmars projectile vomit when threatened (24), which could facilitate the transmission of HPAI within this species or to other seabirds. Future surveillance within these species is necessary to confirm these hypotheses.

One clear result from these studies is that A6 H5N5 viruses are likely under-surveilled. Though our inference of host and geographic patterns are based on all currently available data, transmission in other species and beyond the Atlantic coast could be common for H5N5 viruses, but missing from our dataset due to biases in surveillance efforts. Undersampling of these H5N5 lineages is made clear by the clustering of the Pacific coast sequences (from a human and polar bear) with viruses sampled over 3,000 km away. These two data points suggest at least two instances in which significant, unsampled transmission occurred prior to a mammal infection.

These results highlight the capacity for viruses capable of causing spillovers to circulate undetected for long periods of time, and suggest that enhanced surveillance is necessary for better resolving transmission routes and risk. Enhanced surveillance of these viruses would enable better linking viral transmission patterns with avian ecology and movement, and further resolve paths of possible transmission risk and hosts contributing to spread.

Several features of H5N5 set it apart from H5N1. The Eurasian N5 neuraminidase gene of the A6 lineage contains a deletion not found in other H5N5 viruses, making it somewhat unique (6,7). While H5N1 viruses have repeatedly reassorted with enzootic low pathogenicity strains, H5N5 A6 viruses are entirely Eurasian (6). These patterns could reflect repeated introductions of wholly Eurasian viruses, or could arise if the species supporting H5N5 transmission are less likely to facilitate reassortment. Ecologically, while most H5N1 lineages have spread throughout continental North America via Anseriformes species (4), transmission of the A6 lineage of H5N5 is dominated by seabirds and localized to high latitudes and coast lines. Seabirds and shorebirds have been shown to harbor somewhat distinct lineages of H5N1 in Europe and North America, and are the primary hosts for H13 and H16 low pathogenicity viruses (26), supporting the tendency for distinct viruses to circulate in these hosts. The prevalence of H5N5 in seabirds may also increase exposure risk to marine mammals, which in turn suggests an increased risk for mammalian adaptation. Transmission of avian influenza to marine mammals may occur due to close interactions with seabirds in harbors, at fishing docks, or while feeding out at sea, and has notably caused a large outbreak of HPAI H5N1 in South American sea lions in 2023 (27,28). Seabird associated viruses therefore continue to pose threats to wild marine mammals, and merit continued monitoring. Finally, while H5N1 viruses have caused extensive spillovers into domestic poultry, H5N5 viruses appear to spill over less frequently. Within our dataset, only a single North American Galliformes sequence is available, suggesting far more limited impacts of these viruses on agriculture. Whether this pattern arises because these viruses cause fewer spillovers, cause less severe disease, or both, remains unknown.

The dataset used in this paper is a comprehensive collection of sequences from around the world. However, limitations remain. Ecology and animal movement are immensely difficult to accurately model, and it remains possible that the true ecological story is far more complex. Of particular note is the possibility of transmission within avian hosts that did not show clinical signs of HPAI infection, and therefore were not sampled. Additionally, this paper uses a relatively small dataset, and sampling is biased towards North America—in particular, Massachusetts—so this region may appear disproportionately important to transmission. Finally, the long branches seen on parts of the phylodynamic tree indicate periods of cryptic transmission, when H5N5 viruses were circulating through hosts without regular sampling. This is particularly concerning given the long branches associated with movement up and across North America into the Pacific Northwest, and indicates that these viruses are likely under-sampled.

Since its introduction into North America, surveillance and monitoring efforts for HPAI viruses are focused on H5N1 viruses in Anseriformes species. However, our data show that H5N5 viruses have been repeatedly introduced and have persistently co-circulated alongside predominant H5N1 strains, but with wholly distinct ecologies. These H5N5 viruses exhibit host distributions and geographic patterns of spread that are markedly different from those of H5N1, necessitating alternative surveillance strategies that are formulated to these lineages. The close link between H5N5 transmission and seabird and shorebird ecology suggests that these viruses may continue to move across continents and circulate cryptically in these species, while the human infection shows that these viruses could pose spillover risk. Increased surveillance in wild birds and mammals, as well as monitoring of key locations of repeated viral introductions, would be crucial for the prevention of further spread amongst wild birds and for the prevention of potential spillover into mammals.

## Methods

### Surveillance and sample collection

Samples were obtained from both live and deceased animals through collaborative efforts with wildlife clinics and rehabilitators, local, state and federal partnerships, and carcass sampling from mortality events, with each facility adhering to their institution or agency animal handling and permitting procedures. The study was approved by the Tufts University Institutional Animal Care and Use Committee (IACUC) under protocol G2023-90. Most animals were sampled by pooled oropharyngeal and cloacal swabs; when feasible a brain swab was also collected from carcasses. Samples were collected on flocked polyester tipped swabs (Puritan Medical Products, Guilford, ME, USA) placed into viral transport media (VTM) comprised of Medium 199, nystatin, gentamicin, benzylpenicillin, streptomycin, sulfamethoxazole, kanamycin sulfate (Sigma-Aldrich, St. Louis, MO) and bovine serum albumin (ThermoFisher Scientific, Waltham, MA, USA). Samples were stored at -80°C or -20°C until processed.

### Viral sequencing and consensus sequence generation

RNA was extracted from 50µl of viral transport media (VTM) per sample using the Mag-Bind Viral DNA/RNA 96 kit (Omega Bio-Tek Inc., Norcross, GA, USA) on a semiautomated KingFisher Purification System robot (ThermoFisher Scientific, Waltham, MA, USA). RNA was screened for the influenza A virus (IAV) Matrix (MP) gene by reverse transcription polymerase chain reaction (RT-PCR) on the StepOnePlus platform (ABI, Beverly, MA). 5μl total RNA was added to qScript XLT One-Step RT-qPCR ToughMix ROX (VWR, Franklin, MA) with forward M F25 (5’-AGATGAGTCTTCTAACCGAGGTCG-3’), reverse 2002 M R124 (5’- TGCAAAAACATCTTCAAGTCTCTG-3’) and probe M P64 (FAM-TCAGGCCCCCTCAAAGCCGA-TAM) oligonucleotides and a one-step real time RT-PCR run was performed. Any sample with a Ct <40 was further screened for hemagglutinin H5 using the same protocol described above but with forward H5.2344-1673Forward (5’- TACCAAATAYTGTCAATTTATTCAAC-3’), H5.2344-1749Reverse (5’- GTAAYGACCCRTTRGARCACATCC-3’) and probe H5.2344-1718Probe (FAM-CTGGCAATCATDRTGGCTGGTCT-TAM) oligonucleotides. Samples with a Ct <35 were sent to the National Veterinary Services Laboratories for further confirmatory RT-PCR testing and sequencing. A subset of samples were sequenced at the University of Michigan. The 8 genomic segments of influenza A virus were amplified in a one-step multiplex reverse transcription polymerase chain reaction (RT-PCR) using universal IAV primers, uni12/inf1 forward (5’- GGGGGGAGCAAAAGCAGG-3’), uni12/inf3 forward (5’-GGGGGGAGCGAAAGCAGG-3’) and uni13/inf1 reverse (5’-CGGGTTATTAGTAGAAACAAGG-3’) using the SuperScript™ III One-Step RT-PCR System with Platinum™ Taq High Fidelity DNA Polymerase (ThermoFisher Scientific, Waltham, MA, USA). Amplified nucleic acid was prepared for Oxford Nanopore Sequencing using the Native Barcoding Kit 96 V14 (Oxford Nanopore Technologies, Oxford, United Kingdom) and sequenced on a R10 flow cell on an Oxford Nanopore Technologies GridION MK1. Sequence reads were aligned to the nearest influenza A virus reference and sample consensus sequences were generated using IRMA (29). Influenza clades were assigned by submitting consensus hemagglutinin sequences to Nextclade (30) (https://github.com/moncla-lab/h5-nextclade) using the H5Nx clade 2.3.4.4 reference option. Samples that had all eight segments with ≥90% coverage were submitted to NCBI GenBank (31).

### Sequence data curation

A global H5 clade 2.3.4.4b A6 genotype dataset was compiled by querying the Global Initiative on Sharing All Influenza Data (GISAID) and NCBI Virus (GenBank) databases (10,11). The GISAID search included all H5 clade 2.3.4.4b samples classified as A6 with available genomic sequences for all eight gene segments (accessed on December 16, 2025). The NCBI Virus query was conducted by searching for all H5N5 viruses collected from November 1, 2021 to December 16, 2025 (accessed on December 16, 2025); only samples with available sequences for all eight gene segments were retained. These sequences were then combined with sequences from the samples described above, collected by Tufts. Samples were then assigned a genotype using the GenoFLU genotyping tool (32) (https://github.com/moncla-lab/GenoFLU-multi) to confirm the A6 genotype classification for GISAID sequences and compile an A6-only dataset from the NCBI Virus dataset. The A6 datasets were then merged and de-duplicated based on the isolate ID for each sample and cross referenced to reflect a single sample per animal. An initial dataset of n = 342 whole genomes was generated, with additional genomes added from the University of Michigan in March 2026 and from GISAID (10) and NCBI Virus (11) in May 2026, for a final dataset of n = 355 HPAI H5N5 genomes. Each gene sequence dataset was aligned using MAFFT v.7.526 (33) and trimmed to the coding sequence region. Because GenoFLU genotypes represent non-reassorting groups of sequences, we concatenated all A6 genomes for analyses to enhance phylogenetic inference (34). All eight segment alignments were concatenated using ConcatenatorGUI (35) (https://github.com/iTaxoTools/ConcatenatorGui). One genome (A/american crow/USA/003298-003/2025) was removed from all analyses due to a lack of geographic metadata, for a final count of n = 354 genomes.

### Annotation of discrete and continuous traits

We annotated each sequence in our dataset with additional metadata on geographic location and host. We assigned the continent of samples as a discrete location (described as “region” in figures) for samples from Europe (including Greenland) and Asia (Japan only). Samples from North America were divided into “eastern Canada” (samples from Ontario, Newfoundland, Nova Scotia, and Prince Edward Island Provinces), “northern Canada” (samples from Nunavut Province), “northeastern United States” (samples from Maine, Massachusetts, and Ohio), or “northwestern United States” (Alaska and Washington). Samples from Europe and Asia were assigned the latitude and longitude of the centroid of the country of origin reported on Apple maps or Google Earth as the geographic coordinates, while samples from North America were assigned the finest scale geographic coordinates possible (6 samples from North America had state or province level coordinates, 160 had county, town, or the equivalent as the coordinates).

Host group assignments were based on host taxonomic order and ecological behavior. Ducks, geese, eiders, mergansers, shelducks, and buffleheads were grouped as “goose, duck, seaduck”. Hawks, kites, eagles, owls, and kestrels were grouped as “bird-of-prey”. Chickens, turkeys, and pheasants were grouped as “ground-birds”. While corvids are within the Passeriformes order, we grouped them as “scavengers” with vultures, due to shared behavior and a shared potential transmission pathway. Gulls, terns, kittiwakes, cormorants, fulmars, gannets, skuas, and sandpipers were grouped as “seabirds”. All mammals, both human and non-human, were grouped as “mammal”. The lone environmental sample was simply assigned a discrete host trait of “other”.

### Phylodynamic analysis of discrete and continuous traits

To assess the evolution, movement, and introduction of H5N5 A6 viruses into and throughout North America, we simultaneously conducted a Bayesian continuous phylogeographic analysis and discrete trait analysis with Markov jumps and rewards. We first generated empirical trees using BEAST X v10.5.0 (36). We modeled H5N5 A6 evolution using a HKY + Γ4 substitution model, an uncorrelated relaxed clock model with a mean of 0.005, and any uncertain dates sampled from a period of one year. We tested three models of demographic growth: a constant population model, a model of exponential growth, and a GMRF Bayesian SkyRide model. All three models were run for 300,000,000 MCMC states in BEAST X v10.5.0 (36) for three independent iterations, and combined after removal of 10% burnin to achieve an Effective Sample Size >200 in Tracer v1.7.2 (37). Model fit was assessed using Stepping Stone sampling of 100 steps, with the exponential growth model having the highest marginal likelihood estimation (12). Trees and log files were combined using the LogCombiner v10.5.0 software, and 10% burn-in was removed. Based on these results, an exponential growth model was inferred as the best fit for the data. All subsequent analyses were performed with this exponential model.

We next sampled 867 trees from the posterior distribution of tree topologies and used these as empirical trees for inference of continuous phylogeography (38) and discrete trait reconstruction (39). The continuous phylogeography was inferred using a Cauchy relaxed random walk model with 0.01 jitter for continuous phylogeographic traits. We defined two discrete traits as described above, for host and geography. For each, we performed Bayesian stochastic search variable selection (BSSVS), and mapped transitions using Markov Jumps and Rewards (39). Trait runs were run for 200,000,000 MCMC states in BEAST X v10.5.0 (36) for three independent iterations, and combined after removal of 10% burnin to achieve an Effective Sample Size >200 in Tracer v1.7.2 (37). Trees and log files were combined using the LogCombiner v10.5.0 software, and Maximum Clade Credibility trees were generated with TreeAnnotator v10.5.0. Bayes factors were calculated using spreaD3 v.0.9.7.1 (40), and geographic dispersal maps were generated using SERAPHIM 2.0 (41). Trees were visualized with Baltic (42). All analyses of Markov jumps and dispersal statistics (i.e. weighted diffusion coefficient and distance from introduction point) were performed on a posterior distribution of 834 trees. Markov reward proportions were performed on the posterior distribution of 834 trees, but Markov rewards over time were computed from the Maximum Clade Credibility. Trees visualized in Baltic (42) and maps generated with SERAPHIM (41) were built using the Maximum Clade Credibility tree.

## Data availability

All code used to generate the results and analyses are publicly available at https://github.com/moncla-lab/H5N5-seabird-phylo.git. All sequences used in this analysis are available as an EPI set from GISAID under EPI_SET_261001ep and are acknowledged in the Supplementary Appendix.

## Acknowledgments

We gratefully acknowledge all data contributors, i.e., the Authors and their Originating laboratories responsible for obtaining the specimens, and their Submitting laboratories for generating the genetic sequence and metadata and sharing via the GISAID Initiative, on which this research is based. We would like to thank Allison Black and Hanna Oltean for their feedback and comments on the manuscript. This work was supported by the National Institute of Allergy and Infectious Diseases, National Institutes of Health, Department of Health and Human Services (co. number 75N93021C00016) and by grant number R01AI195533. Viral sequencing in the Lauring laboratory was supported by U19AI181767. Clara Malekshahi was supported by funding from NIH Grant 1F31AI202942-01. This research was supported in part by the Intramural Research Program of the National Institutes of Health (NIH). The contributions of the NIH author(s) are considered Works of the United States Government. The findings and conclusions presented in this paper are those of the author(s) and do not necessarily reflect the views of the NIH or the U.S. Department of Health and Human Services, or the United States government. This work was conducted with support from the Research Corporation for Science Advancement (RCSA) as part of Subaward No. SA-MZT-2023-065b, part of a USDA Non-Assistance Cooperative Agreement with RCSA Federal Award No. 58-3022-0-005. USDA is an equal opportunity provider and employer.

## Author Contributions

W.P., M.I.N., N.S.C.K., and L.H.M. conceived the project. N.S.C.K., A.S., and W.P. collected samples and produced data. S.G. and A.L.G. sequenced the virus isolated from a human in Washington state. L.P., W.J.F., and A.S.L. sequenced eight samples provided by N.S.C.K. and W.P.. A.C.B. and C.R.M. curated data. C.R.M. and L.H.M. designed the experiments. C.R.M. performed the analyses and generated figures. C.R.M., L.H.M., N.S.C.K., A.C.B., W.P., L.P., and W.J.F wrote the manuscript text, and all authors contributed to editing. All authors discussed and approved of the manuscript.

