## Supplementary Figure for "H5N5 virus transmission is driven by repeated cross-Atlantic introductions and transmission among high latitude seabirds"

**Supplementary Figure 1.**

**Supplementary Table 1.**

**Supplementary Figure 2.**

**Supplementary Figure 4.**

**Supplementary Figure 5.**

**Supplementary Figure 6.**

**Supplementary Figure 7.**

**Supplementary Figure 8.**

**Supplementary Appendix.**

**Supplementary Figure 1. Sequences are primarily from samples from Europe and east Canada.** Plot of number H5N5 genomes per month across the study period, colored by region.

| **Supplementary Table 1. Demographic model fit.** Average marginal likelihood estimation scores for three identical interactions of each population growth model. | | |
| --- | --- | --- |
| **Model** | **Stepping Stone Sampling Score** | **Path Sampling Score** |
| Constant Population Size | -45,848 | -45,843 |
| Exponential Growth | -45,830 | -45,825 |
| GMRF SkyRide | -45,850 | -45,843 |

**Supplementary Figure 2. A6 HPAI H5N5 viruses were introduced from Europe to North America at least nine times.** Phylogenetic tree of n = 354 concatenated genomes, colored by known or inferred region, exploded by cross-continental transitions, with origin continent indicated by shape (square = Europe, triangle = North America).

**Supplementary Figure 3. The majority of transmission occurred in Europe and eastern Canada.** Proportion of Markov Rewards for each region across the trunk of the phylogenetic tree.

**Supplementary Figure 4. The timing of transitions between Europe and North America exhibit high variability.** The frequency of transitions from Europe to North America (green) and North America to Europe (blue) over time across the posterior distribution of trees **(A)**. Distribution of transitions from Europe to North America (green) and North America to Europe (blue) per month across the posterior distribution of trees, with a boxplot overlaid (medians in bold) **(B)**.

**A**

**B**

**Supplementary Figure 5. Statistical analysis of transitions between Europe and North America per month.** Poisson regression test of transitions per month from Europe to North America (intercept month = January) **(A)**. Distribution of transitions from Europe to North America over the posterior distribution of trees per month **(B)**. Poisson regression test of transitions per month from North America to Europe (intercept month = January) **(C)**. Distribution of transitions from North America to Europe over the posterior distribution of trees per month **(D)**.

**A**

**B**

**C**

**D**

| **Predictor** | **Coefficient** | **95% Interval** | **p-value** |
| --- | --- | --- | --- |
| Intercept | -0.5702 | -0.660, -0.480 | 0.000 |
| February | 0.0127 | -0.115, 0.140 | 0.845 |
| March | 0.4665 | 0.351, 0.582 | 0.000 |
| April | 0.5785 | 0.466, 0.691 | 0.000 |
| May | 0.3082 | 0.189, 0.427 | 0.000 |
| June | 0.1713 | 0.049, 0.294 | 0.006 |
| July | 0.0557 | -0.070, 0.182 | 0.386 |
| August | -0.0390 | -0.168, 0.090 | 0.554 |
| September | 0.1161 | -0.008, 0.240 | 0.067 |
| October | 0.8707 | 0.763, 0.978 | 0.000 |
| November | 0.9951 | 0.889, 1.101 | 0.000 |
| December | -0.1684 | -0.302, -0.035 | 0.013 |

| **Predictor** | **Coefficient** | **95% Interval** | **p-value** |
| --- | --- | --- | --- |
| Intercept | -0.8728 | -0.978, -0.768 | 0.000 |
| February | -0.1221 | -0.275, 0.031 | 0.119 |
| March | -0.5051 | -0.676, -0.334 | 0.000 |
| April | -1.1427 | -1.356, -0.929 | 0.000 |
| May | -0.9619 | -1.162, -0.762 | 0.000 |
| June | -0.9106 | -1.107, -0.714 | 0.000 |
| July | -0.4185 | -0.585, -0.252 | 0.000 |
| August | 0.1664 | 0.024, 0.309 | 0.022 |
| September | 0.1036 | -0.041, 0.249 | 0.161 |
| October | 0.1267 | -0.017, 0.271 | 0.085 |
| November | 0.2046 | 0.063, 0.346 | 0.005 |
| December | 0.2747 | 0.135, 0.414 | 0.000 |

**Supplementary Figure 6. Global transmission of A6 HPAI H5N5 virus is dominated by seabirds and shorebirds.** Phylodynamic tree of H5N5 A6 viruses, colored by host group **(A)**. Host group probability across all internal nodes of the Maximum Clade Credibilty tree over time, where over 86% of internal nodes infer the host group with over 95% confidence **(B)**. Net Markov transitions (jumps from – jumps to) for each host; host groups below zero are primarily sinks, host groups above zero are primarily sources **(C)**.

**A**

**B**

**C**

**Supplementary Figure 7. Geographic dissemination of HPAI H5N5 A6 viruses is limited to northern latitudes.** Geographic map of Europe and North America showing the movement of inferred (circles) and sequenced (squares) viruses over time (not shown: samples from Russia and Asia).

**Supplementary Figure 8. The clock rate of viruses sampled in the northwestern United States is consistent with the phylogenetic tree as a whole.** Analysis from TempEst showing the root-to-tip divergence over time of the entire phylogenetic tree, with sequences from the northwestern United States indicated.
